# VAMP2 regulates phase separation of alpha-synuclein

**DOI:** 10.1101/2023.06.16.545277

**Authors:** Aishwarya Agarwal, Farheen Raza, Christine Hilcenko, Katherine Stott, Nobuhiro Morone, Alan J. Warren, Janin Lautenschläger

## Abstract

**Summary:** Alpha-synuclein (aSYN) is a small synaptic protein that is linked to Parkinson’s disease (PD) and other synucleinopathies. aSYN has been shown to undergo protein phase separation. We found that aSYN phase separation, in vitro and in cells, is regulated by vesicle-associated membrane protein 2 (VAMP2), the vesicular SNARE protein. We show that aSYN phase separation is promoted by C-terminal electrostatic interaction, which in cells is mediated via interaction with VAMP2. The condensate formation is specific for the R-SNARE VAMP2 and critically dependent on aSYN’s capacity to bind to lipid membranes. In vitro, the addition of VAMP2 decreases the saturation concentration of aSYN phase separation, which is mediated by its juxtamembrane domain. Our results delineate a molecular mechanism for the regulation of aSYN phase separation, indicating a potential switch from the dispersed to the phase-separated state during vesicle cycling.

**Highlights:**

- Alpha-synuclein phase separation is regulated by electrostatic interactions
- VAMP2 initiates alpha-synuclein condensation in cells
- Positive charges within the juxtamembrane domain of VAMP2 initiate alpha-synuclein phase separation in vitro
- Alpha-synuclein condensates nucleate on lipid membranes and assemble vesicular structures in cells

## Introduction

Biomolecular condensation, also known as phase separation, describes the demixing of biomolecules into a highly concentrated dense phase and a depleted dilute phase. The highly condensed phase with weak multivalent interaction offers tight regulation, while the absence of any delimiting membrane facilitates the dynamic exchange of components with the environment ^1–4^. By now biomolecular condensates have been implicated in various complex biological processes ranging from signal transduction to microtubule assembly to gene regulation ^5, 6^. A series of recent findings have also indicated a role of protein phase separation in synaptic transmission and synaptic vesicle (SV) trafficking. For instance, when mixed at an equimolar ratio, the scaffold proteins PSD-95, GKAP, Shank, and Homer of the postsynaptic density undergo phase separation at concentrations well below their synaptic concentrations ^7, 8^. On-demand release of SVs from vesicle clusters can be explained by the fluid-like organization via phase separation of synapsin ^9, 10^. RIM and RIM-BP, which are components of the presynaptic active zone, have been shown to undergo phase separation in vitro and to form condensates that can effectively cluster voltage-gated calcium channels ^11, 12^. Furthermore, phase separation of active zone scaffold proteins liprin-α and ELKS-1 is important for recruiting downstream binding partners ^13, 14^. Finally, phase separation of Eps15/Fcho1/2 and dynamin/syndapin1 has been shown to be involved in endocytosis mechanisms ^15, 16^.

The presynaptic protein alpha-synuclein (aSYN), involved in neurodegeneration and linked to synucleinopathies like Parkinson’s disease (PD) and Lewy body dementia ^17^ has been reported to undergo protein phase separation ^18, 19^. Further studies have confirmed these findings demonstrating the formation of early nanoclusters ^20^ and hardening of aSYN condensates ^21, 22^. High aSYN concentrations are reached within aSYN droplets, estimated at around 30 to 40 mM, which is relevant for the transition of condensates into the aggregation state ^22^. It has been shown that salt and ions affect aSYN phase separation ^23–25^ and that aSYN localizes to synapsin condensates in cells ^26^. Furthermore, aSYN has been described to co-condensate with tau ^27, 28^. However, the physiological relevance of aSYN phase separation has not been demonstrated and to date, a clear mechanism on how aSYN phase separation might be regulated is missing.

aSYN constitutes three main regions, the N-terminal domain, the non-amyloid beta (NAC)-region, and the negatively charged C-terminus (Figure 1A). The N-terminal domain mediates lipid binding forming an amphipathic alpha-helix upon interaction with lipid vesicles ^29–37^. N-terminal residues 6-25 anchor aSYN to the membrane, while residues 26-97 modulate the strength of its lipid interaction ^38, 39^. The hydrophobic NAC region has been implicated in self-association and protein aggregation ^40–42^, while the C-terminal region is intrinsically disordered ^18^ and is neither involved in helix formation ^31–36^ nor in the formation of aSYN fibrils ^43–46^. However, the C-terminal region of aSYN has been shown to interact with synaptic vesicles in the presence of calcium and to influence aSYN localization in synaptosomes ^47^. aSYN can bind to multiple protein partners ^48^ and has been found to interact with the vesicle fusion machinery, in particular synaptobrevin-2 / vesicle-associated membrane protein 2 (VAMP2), the vesicular R-SNARE protein ^49, 50^. In this context, aSYN regulates SNARE-complex assembly ^49, 51, 52^ and SV clustering ^53–55^.

**Figure 1.**
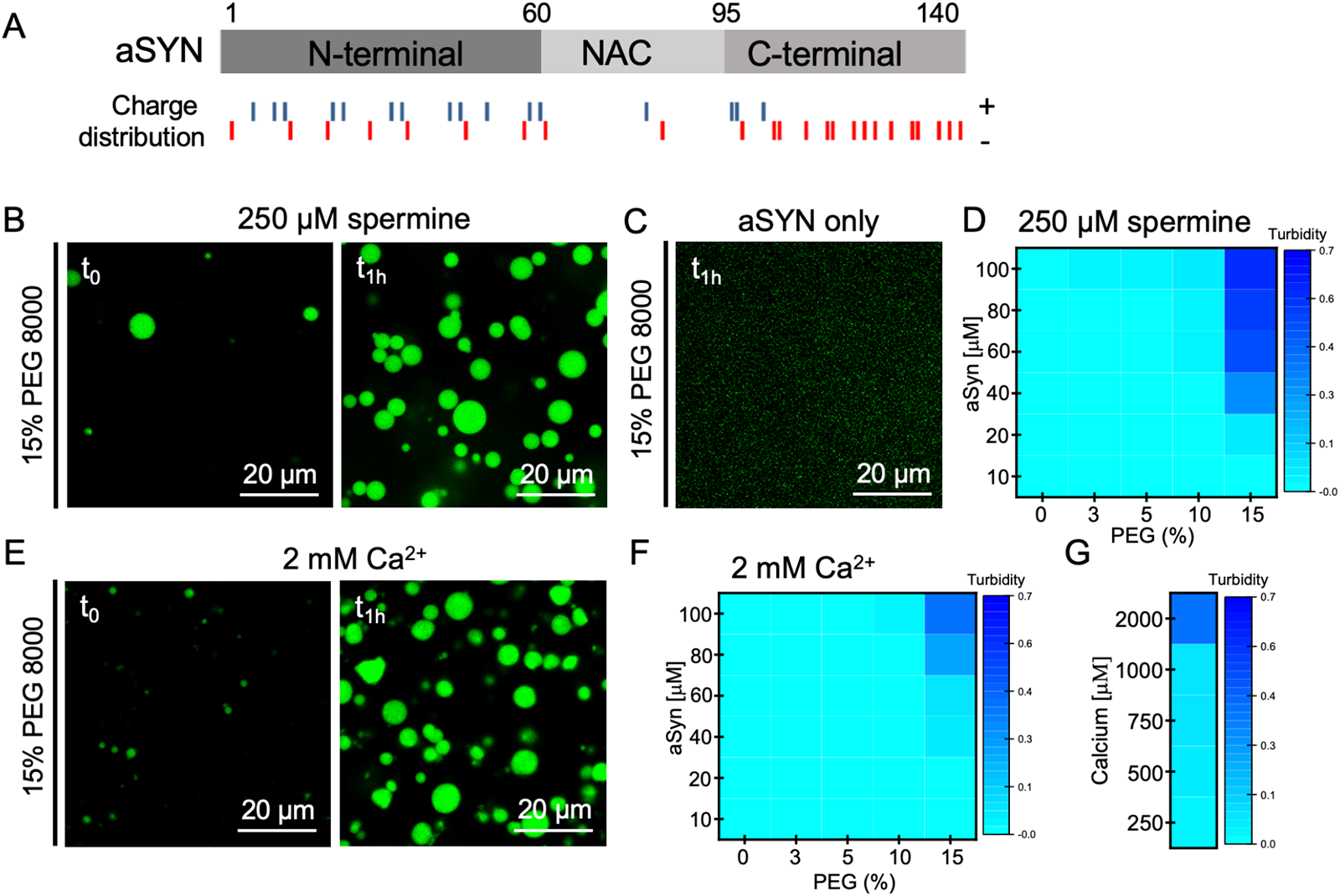
aSYN undergoes phase separation upon electrostatic interaction. (A) aSYN showing its 3 main protein regions. Charge distribution along aSYN sequence; blue – positively charged residues, red – negatively charged residues. (B) aSYN phase separation in the presence of 250 µM spermine and crowding with 15% PEG 8000, immediately after PEG addition (t_0_), and after 1 hour (t_1h_). aSYN concentration used: 100 µM. (C) aSYN on its own does not show droplet formation in the presence of 15% PEG 8000. aSYN concentration used: 100 µM, after 1 hour (t_1h_). (D) Heatmap showing turbidity measurements of aSYN phase separation in the presence of 250 µM spermine. Data represent 3 biological repeats. (E) aSYN phase separation in the presence of 2 mM Ca^2+^ and crowding with 15% PEG 8000, immediately after PEG addition (t_0_), and after 1 hour (t_1h_). aSYN concentration used: 100 µM. (F) Heatmap showing turbidity measurements of aSYN phase separation in the presence of 2 mM Ca^2+^. Data represent 3 biological repeats. (G) Heatmap for aSYN phase separation in the presence of different Ca^2+^ concentrations in the presence of 15% PEG 8000. aSYN concentration used: 100 µM.

In this paper, we demonstrate that VAMP2 is involved in the regulation of aSYN phase separation. We first identified that electrostatic interactions at aSYN’s C-terminal region modulate aSYN phase separation. We then performed a screen for potential interaction partners and found that VAMP2, which interacts with aSYN’s C-terminus ^49, 50^, induces aSYN condensate formation in cells. VAMP2, but not the Q-SNARE’s syntaxin-1A or SNAP25, promotes aSYN condensate formation. Using small synthetic peptides of VAMP2, we further show that positive charges within the juxtamembrane (JM) domain of VAMP2 promote phase separation of aSYN. Finally, we show that aSYN condensate formation is dependent on aSYN’s capacity to bind to lipid membranes and that aSYN condensates accumulate vesicular structures in cells. Our results support the role of aSYN phase separation during vesicle cycling, regulated by the R-SNARE VAMP2.

## RESULTS

### aSYN undergoes phase separation upon electrostatic interaction

aSYN has been studied for protein aggregation, where divalent cations, such as Cu^2+^ and Ca^2+^ ^47, 56–58^, but also the interference with long-range interactions between its N, NAC and C-terminus have been reported to enhance aggregation ^59, 60^. To assess whether long-range interactions and electrostatic interactions play a role in aSYN phase separation we tested the potential contribution of spermine and Ca^2+^ on aSYN phase separation. Spermine, a polyamine with four positive charges, which has previously been shown to bind to aSYN’s negatively charged C-terminus ^61^ and to break aSYN long-range interactions ^59^, enabled aSYN to undergo protein phase separation. When aSYN, in the presence of 250 µM spermine, was subjected to crowding mimicked by 15% PEG 8000 rapid droplet formation occurred, with droplet growth over time (Figure 1B). Under the same conditions, omitting spermine, aSYN did not show droplet formation (Figure 1C). To quantitatively asses aSYN phase separation we performed turbidity measurements in the presence of spermine, indicating the crowding conditions and aSYN concentrations at which aSYN undergoes droplet formation (Figure 1D). Next, we performed droplet and turbidity assays in the presence of Ca^2+^. Similarly, Ca^2+^ did enable aSYN to undergo immediate droplet formation when subjected to crowding with 15% PEG 8000, demonstrating droplet growth over time (Figure 1E). This was seen in the presence of high Ca^2+^ concentrations i.e. 2 mM Ca^2+^, however in contrast to spermine, no turbidity increase was observed with 10% PEG 8000 (Figure 1F) or at low Ca^2+^ concentrations in the µM range (Figure 1G). Since no droplet formation can be observed at physiologically relevant Ca^2+^ concentrations which are estimated to be around 200-300 µM Ca^2+^ or even below that during synaptic stimulation ^62, 63^, we conclude that the effect of Ca^2+^ is mainly electrostatic. Furthermore, we see aSYN phase separation only under high crowding conditions, which indicates that further regulatory factors are likely to be involved.

### VAMP2 enables aSYN condensate formation in cells

We next hypothesized that facilitation of aSYN phase separation could occur upon binding of a protein interaction partner. Therefore, we ectopically expressed synaptic proteins which have previously been correlated to Parkinson’s disease, together with aSYN YFP in HeLa cells, evaluating potential condensate formation in live cells. One of the tested proteins, synaptobrevin-2 / VAMP2 induced aSYN condensate formation in cells (Figure 2A). aSYN YFP overexpression on its own indicates a cytosolic-nuclear distribution and no sign of aSYN aggregation, congruent with the literature ^64, 65^. However, when aSYN YFP was co-expressed with VAMP2, cytosolic aSYN clusters were observed in about 5-10% of the transfected cells, which were not seen upon co-expression of YFP with VAMP2 (Figure 2B/C/D).

**Figure 2.**
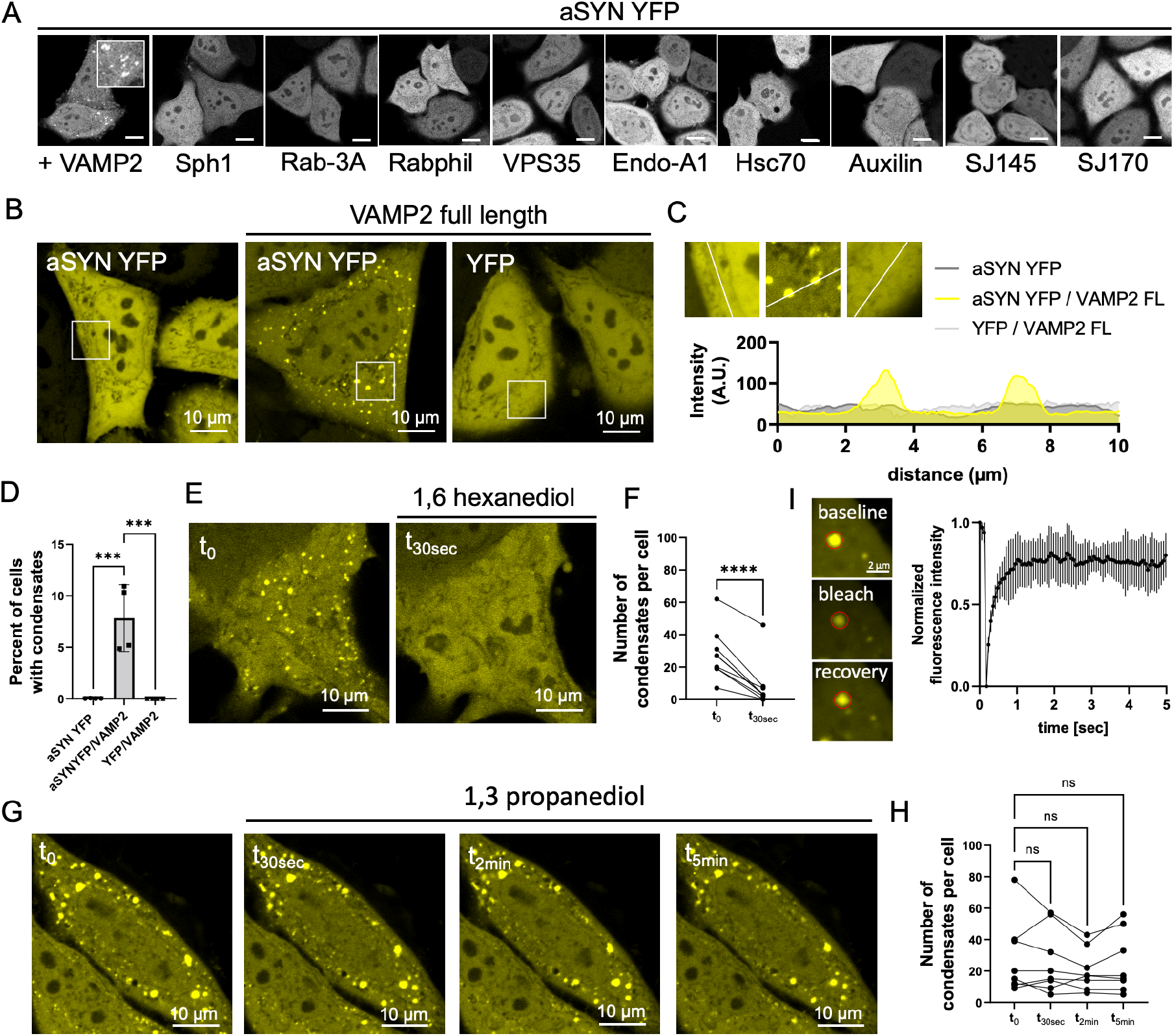
VAMP2 enables aSYN condensate formation in cells. (A) Screening of disease-relevant synaptic proteins on aSYN YFP distribution upon co-expression in HeLa cells. Scale bar 20 µm. VAMP2 – vesicle-associated membrane protein 2; Sph1 – Synphilin-1; Rab-3A – Ras-related protein Rab-3A; Rabphil – Rabphilin-3A; VPS35 – Vacuolar protein sorting-associated protein 35; Endo-A1 – Endophilin-A1; Hsc70 – Heat shock cognate 71 kDa protein; Auxilin – Putative tyrosine-protein phosphatase auxilin; SJ145, SJ170 – synaptojanin-1 isoform 1-145 and 1-170. (B) Cytosolic-nuclear distribution of aSYN YFP upon ectopic expression in HeLa cells, condensate formation upon co-expression of aSYN YFP and VAMP2, co-expression of YFP and VAMP2 shows no condensate formation. (C) Zoom in regions and fluorescence intensity distribution for cells with aSYN YFP only (light grey), aSYN with VAMP2 (yellow), and YFP with VAMP2 (dark grey). (D) Quantification of cells forming condensates. Data derived from incuCyte screening, 16 images per well, 3 wells per biological repeat, 4 biological repeats. Data are represented as mean +/-SD. One-way ANOVA with Dunnett’s multiple comparison test, *p < 0.05; **p < 0.01; ***p < 0.001; ****p < 0.0001. (E) aSYN YFP condensates show dispersal upon incubation with 3% 1,6 hexanediol. See also Figure S1 for recovery of aSYN condensates after 1,6-hexanediol washout. (F) Quantification of condensates per cell, before and after incubation with 3% 1,6-hexanediol. 3 biological repeats, n represents cells. Paired two-tailed t-test, *p < 0.05; **p < 0.01; ***p < 0.001; ****p < 0.0001. (G) aSYN YFP condensates show no dispersal upon incubation with 3% 1,3 propanediol. (H) Quantification of condensates per cell, before and after incubation with 3% 1,3 propanediol. 2 biological repeats, n represents cells. RM one-way ANOVA with Dunnett’s multiple comparison test, *p < 0.05; **p < 0.01; ***p < 0.001; ****p < 0.0001. (I) Photobleaching and recovery of aSYN condensate in cells. Quantification of FRAP experiments. Data are represented as mean +/-SD. 2 biological repeats, n=6.

In order to distinguish the observed clusters from insoluble aggregates we subjected the cells to 1,6-hexanediol, a small aliphatic alcohol, which can dissolve liquid-like assemblies ^66–70^. 3% 1,6-hexanediol caused a rapid dispersal of aSYN clusters as expected for condensates (Figure 2E/F). Furthermore, dissolution of aSYN condensates was reversible after brief wash out periods (Figure S1). 1,3-propanediol, a shorter and therewith more hydrophilic aliphatic alcohol, did not lead to the disassembly of aSYN condensates, reflected by a constant number of condensates per cell after incubation (Figure 2G/H), which indicates a role of hydrophobic interactions within the aSYN condensates ^71^. In addition, fluorescence recovery after photobleaching (FRAP) shows fast recovery times for aSYN condensates with a 0.752 +/-0.066, 0.808 +/-0.004, and 0.831 +/-0.027 recovery after 1, 3 and 5 seconds, demonstrating high mobility between the condensate structures and the cytosolic aSYN fraction (Figure 2I).

These recovery times for aSYN are in line with previous FRAP experiments on synapsin/aSYN co-condensates in cells ^26^. Together, these experiments delineate a liquid-like nature rather than an aggregated state of aSYN assemblies within the cell.

### aSYN/VAMP2 interaction regulates aSYN condensate formation

The vesicular R-SNARE protein, VAMP2 is involved in SNARE complex assembly at the synapse forming a four helical trans-SNARE complex with syntaxin-1A and SNAP-25, which upon vesicle fusion transitions into the cis-SNARE complex, following which VAMP2 is recycled into vesicles ^72–76^. However, VAMP2 has also been shown to be an aSYN interaction partner ^49, 51, 52^, where VAMP2 binding occurs at aSYN’s C-terminal region ^49^. Using alanine scanning of aSYN, the VAMP2 binding site has been mapped to C-terminal residues close to the NAC-region ^50^. Therefore, we ectopically co-expressed aSYN 96AAA, the aSYN mutant with the highest reduction in VAMP2 binding, with VAMP2 in HeLa cells. The aSYN 96AAA mutant was able to form condensates (Figure 3A/B), however, the number of cells forming condensates was significantly reduced (Figure 3C). Furthermore, the size of condensates and the number of condensates per cell decreased (Figure 3D/E). In addition, also the intensity ratio between condensate structures and cytosolic aSYN YFP was reduced, reflecting a lower overall uptake of aSYN into condensates (Figure 3F). To further test that aSYN condensate formation is specific to its interaction with VAMP2, we also co-expressed aSYN YFP with the respective synaptic Q-SNAREs. Both target membrane SNAREs, syntaxin-1A, and SNAP-25, did not promote aSYN condensate formation when ectopically expressed with aSYN YFP in HeLa cells (Figure 3A/B/C).

**Figure 3.**
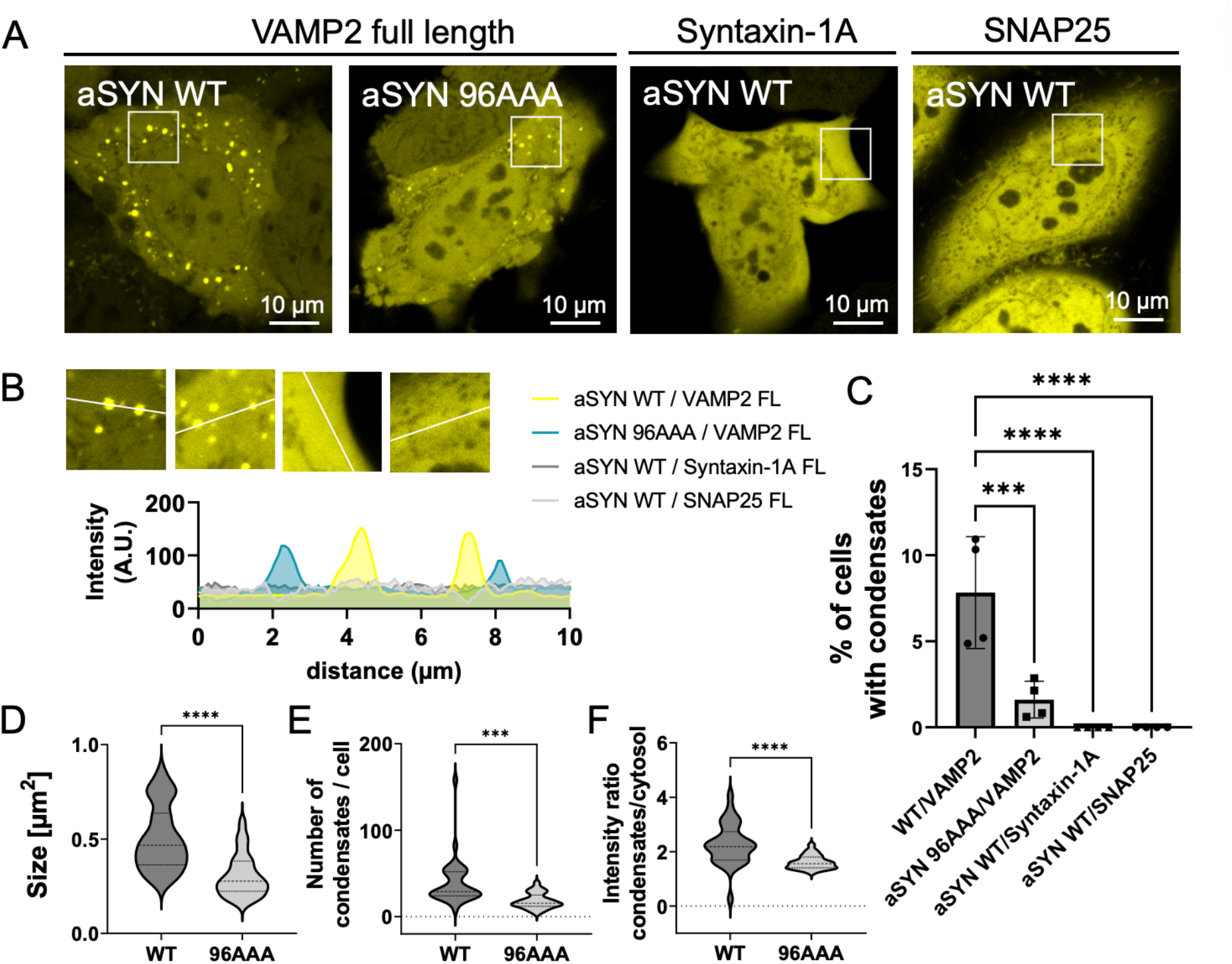
aSYN/VAMP2 interaction regulates aSYN condensate formation. (A) Condensate formation for aSYN wild type (WT) YFP and aSYN 96AAA YFP upon co-expression with VAMP2. aSYN WT YFP co-expressed with syntaxin-1A or SNAP25 does not show condensate formation. (B) Zoom in regions and fluorescence intensity distribution for cells with co-expression of aSYN WT YFP with VAMP2 (yellow), aSYN 96AAA YFP with VAMP2 (turquoise), and aSYN WT YFP with syntaxin-1A (dark grey) and SNAP25 (light grey). (C) Quantification of cells forming condensates. Data derived from incuCyte screening, 16 images per well, 3 wells per biological repeat, 4 biological repeats. Data are represented as mean +/-SD. One-way ANOVA with Dunnett’s multiple comparison test, *p < 0.05; **p < 0.01; ***p < 0.001; ****p < 0.0001. (D) Quantification of condensate size for cells co-expressing aSYN WT YFP and aSYN 96AAA YFP with VAMP2. 3 biological repeats. Data are represented as violin plots. Unpaired two-tailed t-test, *p < 0.05; **p < 0.01; ***p < 0.001; ****p < 0.0001. (E) Quantification of condensates per cell for cells co-expressing aSYN WT YFP and aSYN 96AAA YFP with VAMP2. 3 biological repeats. Data are represented as violin plots. Unpaired two-tailed t-test, *p < 0.05; **p < 0.01; ***p < 0.001; ****p < 0.0001. (F) Quantification of the intensity ratio between condensates and cytosolic aSYN YFP for cells co-expressing aSYN WT YFP and aSYN 96AAA YFP with VAMP2. 3 biological repeats. Data are represented as violin plots. Unpaired two-tailed t-test, *p < 0.05; **p < 0.01; ***p < 0.001; ****p < 0.0001.

### VAMP1-96 promotes aSYN phase separation in vitro

Next, we evaluated whether VAMP2 affects aSYN phase separation in vitro. For our assays, we used VAMP1-96, which, without its transmembrane domain, is a soluble protein. To probe whether VAMP1-96 influences aSYN phase separation we estimated the saturation concentration (C_sat_), the concentration at which aSYN phase separation starts, using a sedimentation-based assay. After induction of aSYN droplet formation upon crowding and centrifugation, the C_sat_ is estimated from the protein level within the supernatant. In the presence of 2 mM Ca^2+^ and 15% PEG 8000 we found a C_sat_ for aSYN phase separation of 33.37 +/-2.36 μM (Figure 4A/B). The addition of VAMP1-96 decreased the C_sat_ of aSYN phase separation to 26.41 +/-1.64 μM (Figure 4A/B). In addition, we observed that VAMP1-96 is pulled down into the pellet fraction when aSYN phase separation is induced indicative of its recruitment to aSYN droplets (Figure 4A lane 4/6, Figure 4C). Therefore, we next probed whether VAMP1-96 would co-condense with aSYN droplets. aSYN phase separation was induced in the presence of 2 mM Ca^2+^ and 15% PEG 8000 as in our previous experiments. The reaction was supplemented with labeled VAMP1-96, demonstrating co-localization of VAMP1-96 with aSYN droplets (Figure 4D/E). As a control, we used the Q-SNARE syntaxin-1A, again as a soluble protein without its transmembrane domain. Labeled Syntaxin1-265 showed significantly less uptake into aSYN droplets (Figure 4D/E), demonstrating that co-localization is specific for VAMP1-96.

**Figure 4.**
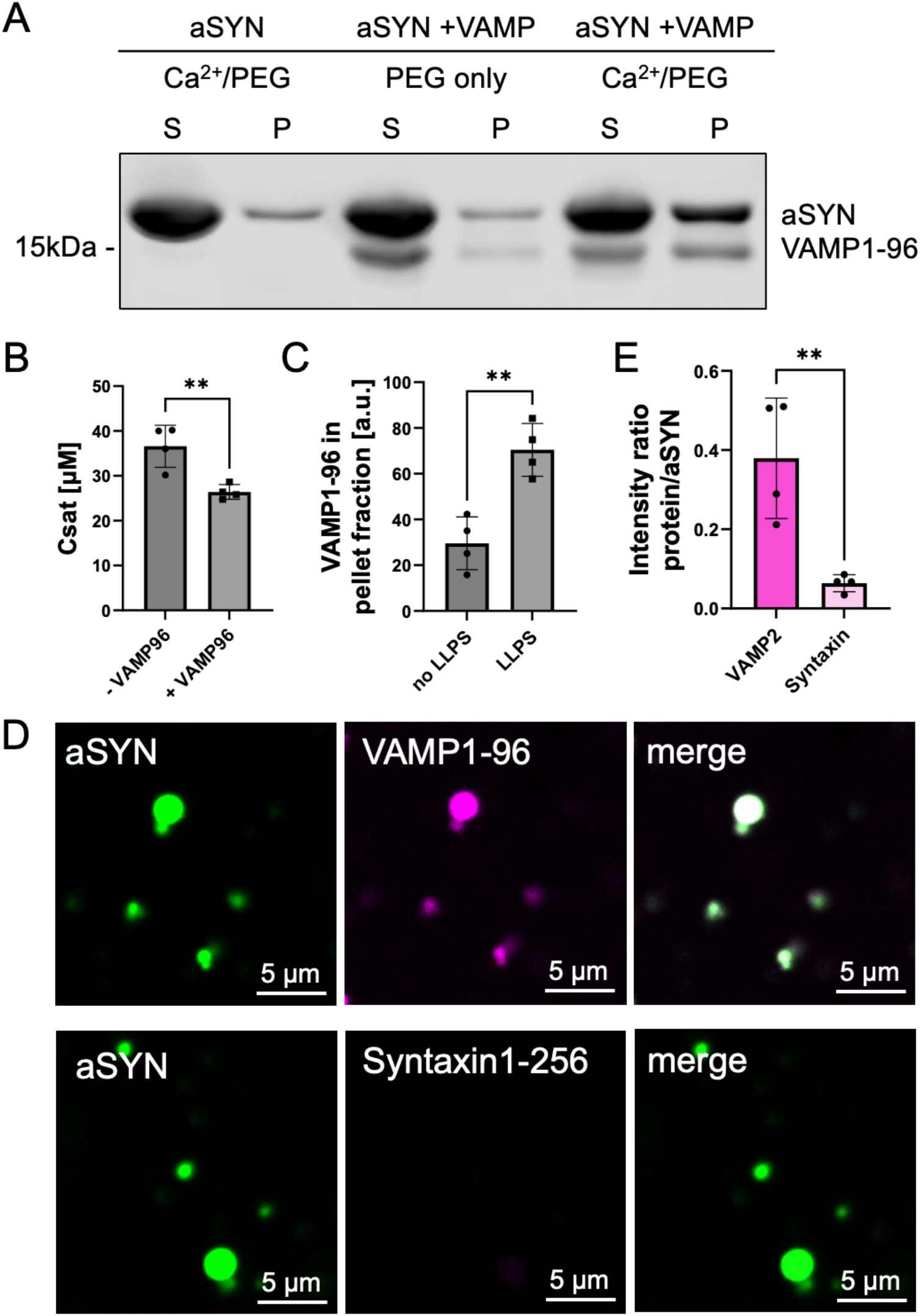
VAMP1-96 promotes aSYN phase separation in vitro. (A) Sedimentation-based assay showing supernatant (S, dilute phase) and pellet (P, droplet) fraction upon aSYN phase separation in the presence of Ca^2+^ (concentrations used: 40 µM aSYN, 2 mM Ca^2+^, 15% PEG), upon aSYN and VAMP1-96 incubation (40 µM aSYN, no Ca^2+^, 10 µM VAMP1-96, 15% PEG) and aSYN phase separation in the presence of VAMP1-96 (40 µM aSYN, 2 mM Ca^2+^, 10 µM VAMP1-96, 15% PEG). (B) Quantification of saturation concentration (C_sat_) of aSYN phase separation in the presence of 2 mM Ca^2+^ (-VAMP) and 2 mM Ca^2+^ with 10 µM VAMP1-96 (+ VAMP). Data from 4 biological repeats. Data are represented as mean +/-SD. Unpaired two-tailed t-test, *p < 0.05; **p < 0.01; ***p < 0.001; ****p < 0.0001. (C) Quantification for the intensity of VAMP1-96 in the pellet fraction, either under no phase separation (40 µM aSYN, no Ca^2+^, 10 µM VAMP1-96, 15% PEG) or under aSYN phase separation conditions in the presence of Ca^2+^ (40 µM aSYN, 2 mM Ca^2+^, 10 µM VAMP1-96, 15% PEG). Data from 4 biological repeats. Data represented as mean +/-SD. Unpaired two-tailed t-test, *p < 0.05; **p < 0.01; ***p < 0.001; ****p < 0.0001. (D) Co-localization of VAMP1-96 or Syntaxin1-265 with aSYN droplets induced in the presence of 2 mM Ca^2^ and 15% PEG 8000. Co-localization was evaluated 30 min after induction of aSYN phase separation and addition of the respective labeled protein. aSYN concentration used: 100 µM. (E) Quantification of co-localization showing intensity ratio of VAMP1-96 and syntaxin1-265 to aSYN after 30 min. Data from 4 biological repeats. Data represented as mean +/-SD. Unpaired two-tailed t-test, *p < 0.05; **p < 0.01; ***p < 0.001; ****p < 0.0001.

### The JM-domain of VAMP2 enables aSYN phase separation

We next tested which region of VAMP2 is involved in inducing aSYN phase separation. We focused on two regions that have been previously described to interact with aSYN, the N-terminal domain of VAMP2 ^49^, but also a more C-terminal region that has been shown to mediate aSYN/VAMP2 binding during aggregation ^77^. We used two peptides, one resembling residues 25-30 of the N-terminal domain, and one resembling residues 83-88 in the JM-domain of VAMP2 (Figure 5A). In the droplet as well as in the turbidity assay the N-terminal peptide (NT peptide, NLTSNR) did not influence aSYN phase separation (Figure 5B/C). However, the JMD peptide (KLKRKY) was found to induce aSYN droplet formation (Figure 5B) and showed increased turbidity (Figure 5C). In order to test specificity and to cover larger protein regions, we designed longer peptides, covering either the first half of the N-terminal domain (NT long peptide 1, aa 3-16, ATAATAPPAAPAGE), the second half of the N-terminal domain (NT long peptide 2, aa 17-30, GGPPAPPNLTSNR), the full length of the JM-domain (JMD long peptide, aa 83-96, KLKRKYWWKNLKMM), or the SNARE/JMD interlinking region including the residues of the previous short JMD peptide (SNARE/JMD long peptide, aa75-88, SQFETSAAKLKRKY, Figure 5D). The droplet assay and the turbidity measurements show that the NT peptides and the SNARE/JMD long peptide do not induce aSYN phase separation, while the JMD long peptide enabled aSYN droplet formation indicating the role of the JM-domain in promoting aSYN condensate formation (Figure 5E/F). Further, to test the sequence specific effect on phase separation we designed a scrambled JMD long peptide (Peptidenexus scrambler, KMLKWKMNKYLRWK). The native JMD peptide showed a significant stronger effect than the scrambled JMD long peptide, demonstrating that beside electrostatic interactions the distribution of charged and uncharged residues within the JM-domain is important to mediate specificity.

**Figure 5.**
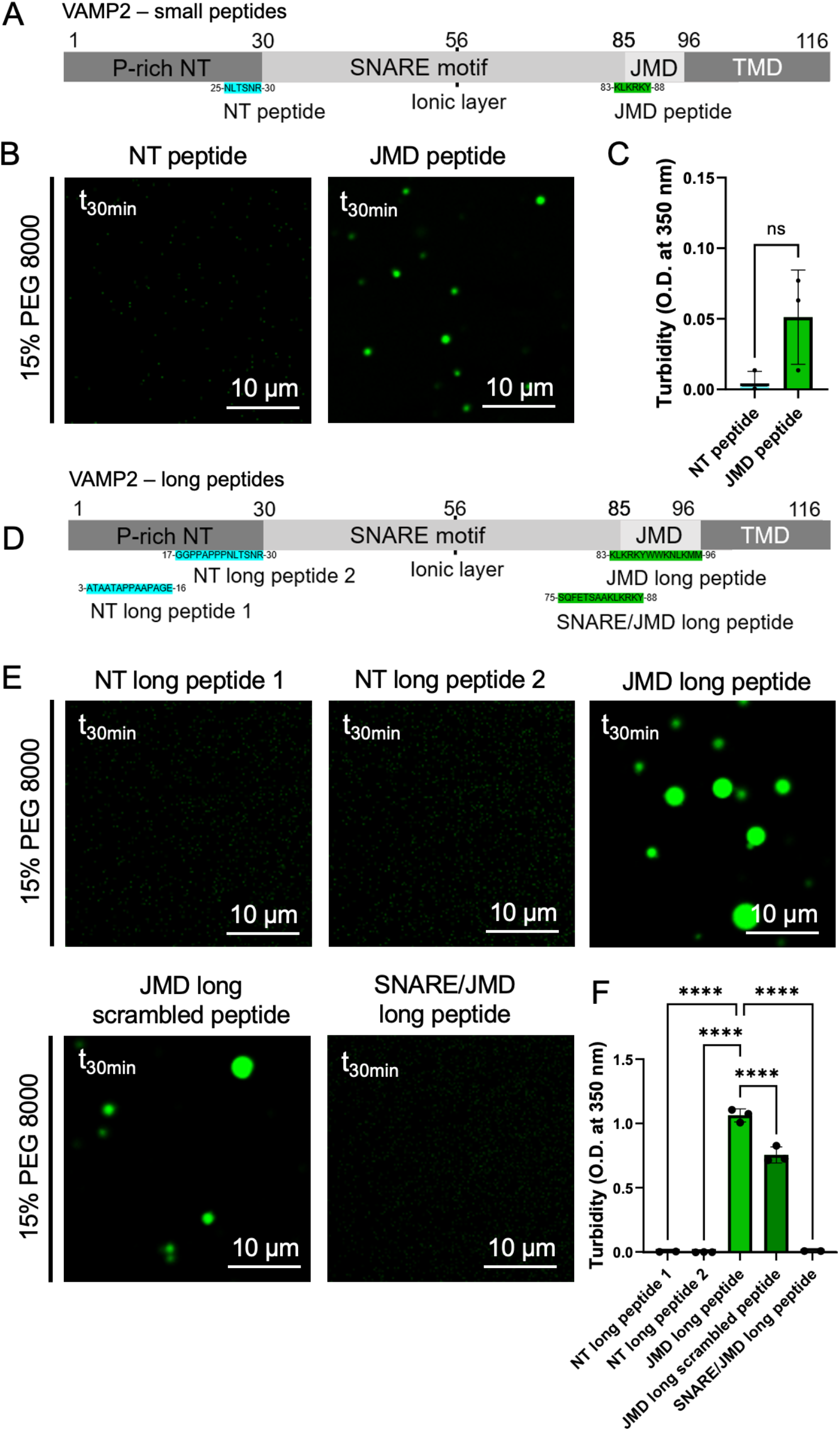
The JM-domain of VAMP2 enables aSYN phase separation. (A) Schematic of VAMP2 with the respective N-terminal peptide (NT peptide) and JM-domain peptide (JMD peptide). (B) Evaluation of aSYN phase separation in the presence of 150 µM peptide and crowding with 15% PEG 8000, 30 min after PEG addition (t_30min_), 40 µM aSYN. (C) Quantification of turbidity measurements at t_30min_. Data are represented as mean +/-SD. 3 biological repeats. Unpaired two-tailed t-test, *p < 0.05; **p < 0.01; ***p < 0.001; ****p < 0.0001. (D) Schematic of VAMP2 with the respective N-terminal long peptides (NT long peptide 1 and 2), the native JMD long peptide, and the SNARE/JMD long peptide. (E) Evaluation of aSYN phase separation in the presence of 150 µM peptide and crowding with 15% PEG 8000, 30 min after PEG addition (t_30min_), 40 µM aSYN. (F) Quantification of turbidity measurements at t_30min_. Data are represented as mean +/-SD. 2-3 biological repeats. One-way ANOVA with Dunnett’s multiple comparison test, *p < 0.05; **p < 0.01; ***p < 0.001; ****p < 0.0001.

### The JM-domain of VAMP2 interacts with aSYN

To further validate the effect of the JMD long peptide we estimated the C_sat_ for aSYN phase separation in the presence of peptide using the sedimentation-based assay as described above. A significant decrease in aSYN C_sat_ was seen for 50 μM JMD long peptide. At 100 μM JMD long peptide a further decrease in aSYN C_sat_ to 11.54 +/-3.46 μM was observed, while concentrations of 150 μM JMD long peptide and above showed a stable reduction to a C_sat_ of about 3 to 4 μM (Figure 6 A/B). Using 150 μM JMD long peptide we next performed turbidity measurements, which demonstrate that aSYN phase separation is induced down to 10 µM aSYN in the presence of 15% PEG 8000 and down to 20 µM aSYN in the presence of 10% PEG 8000 (Figure 6C). Furthermore, we observe droplet formation of aSYN in the presence of JMD long peptide at 10% and 5% PEG 8000 (Figure S2).

**Figure 6.**
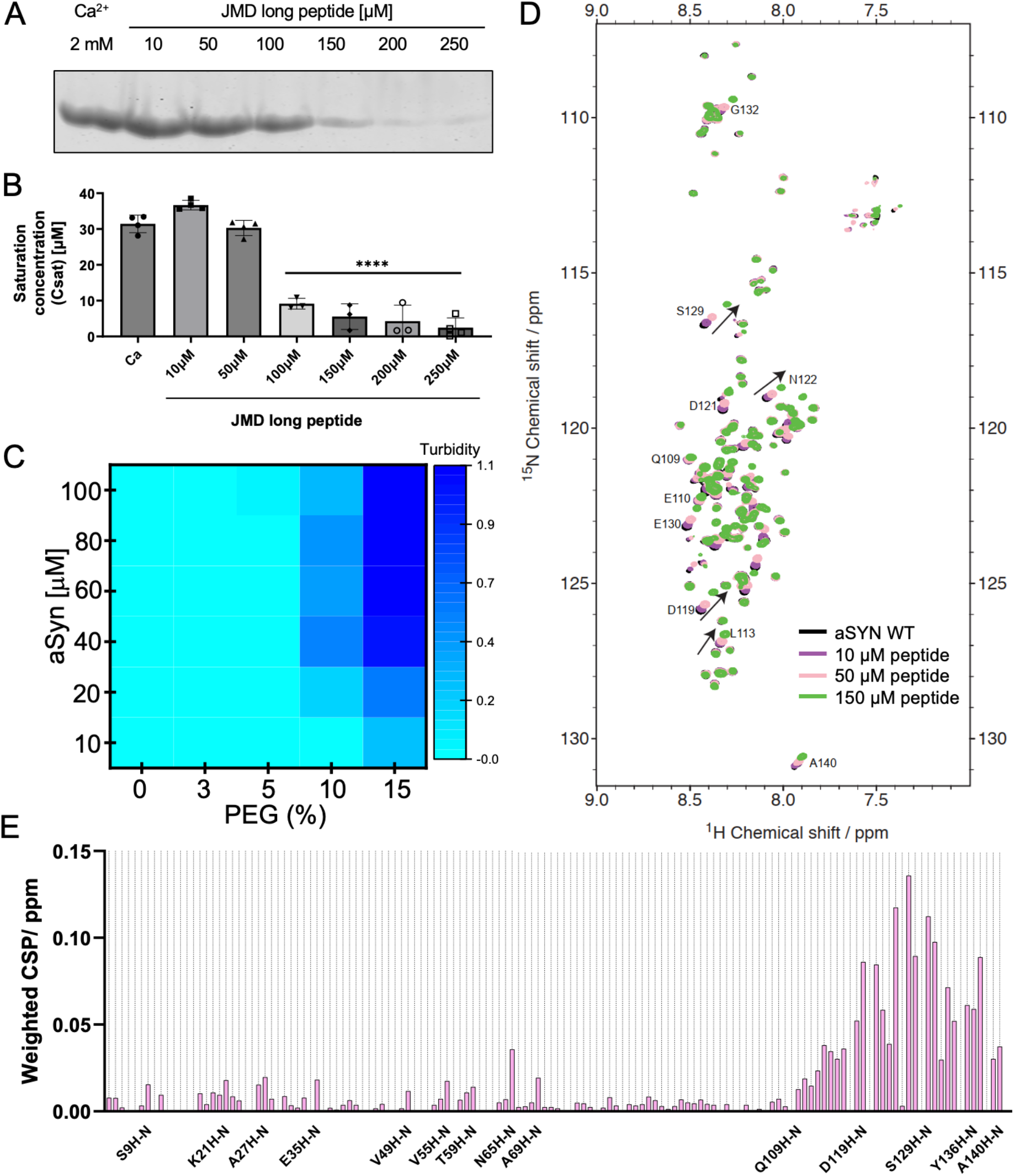
The JM-domain of VAMP2 interacts with aSYN. (A) Sedimentation-based assay showing supernatant (dilute phase) upon aSYN phase separation in the presence of the JMD long peptide (40 µM aSYN, 15% PEG, concentrations of peptide as indicated). (B) Quantification of saturation concentration (C_sat_) of aSYN phase separation in the presence of the JMD long peptide. Data represented as mean +/-SD. n represents biological repeats. One-way ANOVA with Dunnett’s multiple comparison test, *p < 0.05; **p < 0.01; ***p < 0.001; ****p < 0.0001. (C) Heatmap showing turbidity measurements of aSYN phase separation in the presence of 150 µM JMD long peptide. Data represent 3 biological repeats. See also Figure S2 for aSYN droplet formation in the presence of 150 µM JMD long peptide at lower PEG concentrations. (D) Overlapped ^1^H-^15^N-BEST-TROSY spectra of aSYN without (black) and in the presence of increasing concentrations of JMD long peptide as indicated. See also Figure S3A for the ^1^H-^15^N-BEST-TROSY spectra of aSYN in the presence of NT long peptide 1. aSYN concentration used: 40 µM, peptide concentrations: 10 µM, 50 µM, 150 µM. (E) Weighted average (of ^15^N and ^1^H) chemical shift perturbation 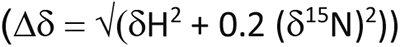 of residues in aSYN in the presence of 50 μM JMD long peptide. Also see Figure S3B weighted average (of ^15^N and ^1^H) chemical shift perturbation in the presence of NT long peptide 1.

In order to test whether and where the JMD long peptide interacts with aSYN we applied multi-dimensional NMR spectroscopy to perform chemical shift perturbation titrations on aSYN using the JMD long peptide concentrations estimated above. The resulting chemical shifts were most prominent at the C-terminal region of aSYN, consistent with an electrostatic interaction. However, smaller chemical shifts were also observed at N-terminal residues and residues within the NAC region (Figure 6D/E). In contrast, the NT long peptide did not induce chemical shifts at the respective residues (Figure S3A/B).

### aSYN condensates initiate on lipid membranes and assemble vesicular structures

Since both, VAMP2 via its transmembrane domain, and aSYN via the formation of an amphipathic alpha-helix bind to lipid membranes, we next hypothesized that aSYN condensate formation occurs on lipid membranes. To test the relevance of lipid binding we used aSYN A30P, an aSYN disease variant ^78^ with defective lipid binding ^30, 37, 79–81^ little effect of A30P. Upon co-expression of aSYN A30P YFP with VAMP2 no condensate formation was observed (Figure 7A/B). These findings suggest that the lipid membrane association of aSYN is important to allow aSYN condensate formation. It is interesting to note that aSYN A30P has reduced but not completely abolished lipid binding, being diminished to about half of aSYN wild-type binding ^80^. Therefore, reduced rather than abolished condensate formation might have been expected. Our results indicate that in cells aSYN A30P might not reach the critical concentration at lipid membranes to nucleate aSYN phase separation. This is consistent with previous findings on the membrane accumulation of aSYN A30P in yeast cells and neurons^82, 83^. aSYN wild-type and aSYN A53T are associated with the plasma membrane when expressed in yeast, while aSYN A30P shows a diffuse cytoplasmic distribution ^82^. Similarly, aSYN A30P behaves like a fully soluble protein upon photobleaching in synaptic terminals, while aSYN wild-type recovers more slowly, indicative of its membrane-bound fraction. Furthermore, upon synaptic stimulation, there is no redistribution seen for aSYN A30P, while aSYN wild-type, similar to synapsin, dissociates upon exocytosis ^83^.

**Figure 7.**
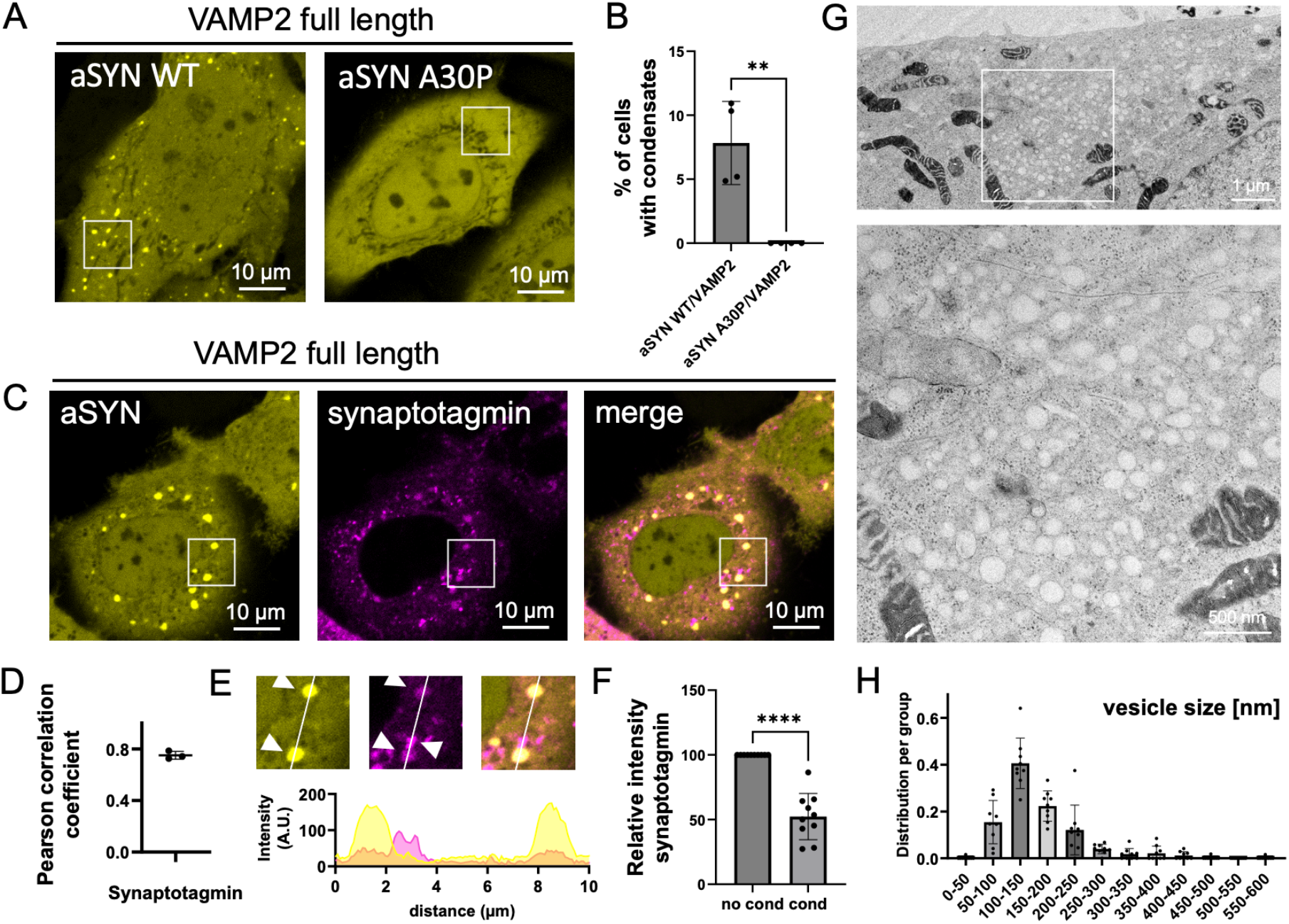
aSYN condensates initiate on lipid membranes and assemble vesicular structures. (A) Co-expression of VAMP2 and aSYN WT YFP in HeLa cells showing condensate formation. Cells upon co-expression of VAMP2 and aSYN A30P YFP, a disease variant with decreased lipid binding, lack condensate formation. (B) Quantification of cells forming condensates. Data derived from incuCyte screening, 16 images per well, 3 wells per biological repeat, 4 biological repeats. Data represented as mean +/-SD. One-way ANOVA with Dunnett’s multiple comparison test, *p < 0.05; **p < 0.01; ***p < 0.001; ****p < 0.0001. (C) Co-expression of aSYN YFP, VAMP2, and mScarlet synaptotagmin. (D) Quantification of Pearson Correlation Coefficient for aSYN YFP and mScarlet synaptotagmin co-localization. 3 biological repeats. Data represented as mean +/-SD. Unpaired two-tailed t-test, *p < 0.05; **p < 0.01; ***p < 0.001; ****p < 0.0001. (E) Zoom in areas and arrowheads highlighting co-localization of aSYN condensates with co-expressed mScarlet synaptotagmin and mScarlet synaptotagmin outside of aSYN condensates. Fluorescence intensity distribution for aSYN YFP (yellow) and mScarlet synaptotagmin (magenta). Enrichment of mScarlet synaptotagmin in aSYN condensates, synaptotagmin positive vesicular structures outside of aSYN condensates with higher intensity. (F) Quantification of mScarlet synaptotagmin intensity. Data from 6 biological repeats. Data represented as mean +/-SD. Unpaired two-tailed t-test, *p < 0.05; **p < 0.01; ***p < 0.001; ****p < 0.0001. (G) Electron microscopy of HeLa cells upon ectopic expression of aSYN YFP and VAMP2 showing assemblies of vesicular structures in the cytosol. (H) Histogram showing the size distribution of vesicular structures. Data represented as mean +/-SD. n represents vesicle clusters analyzed.

Further to these experiments we examined whether aSYN condensates indeed have a regulatory role on vesicle clustering, such as found upon formation of synapsin condensates in cells ^84^. We co-expressed synaptotagmin, a vesicle-bound transmembrane protein involved in Ca^2+^ sensing, together with aSYN YFP and VAMP2. While mScarlet synaptotagmin shows small vesicular structures on its own, it also demonstrates co-localization with the larger aSYN condensates (Figure 7C/D). Further to that, we find that synaptotagmin within condensates shows lower fluorescence intensity compared to vesicular structures outside of condensates (Figure 7E), indicating a secondary uptake of vesicles. Moreover, evaluated using electron microscopy, cells co-expressing aSYN YFP and VAMP2 demonstrate cytosolic accumulation of vesicles (Figure 7F). The observed vesicle clusters harbor vesicles with a wide distribution of sizes, with the main proportion being vesicles between 100-150 nm (Figure 7G). This indicates that aSYN serves a distinct vesicle clustering mechanism, since synapsin condensates do cluster smaller vesicles ^84, 85^.

## DISCUSSION

With the concept of protein phase separation new perspectives for our understanding of cell compartmentalization and on how intracellular processes are organized in space and time emerge ^86–88^. This is true for cytosolic and nuclear phase separated compartments, but also for the organization within the presynaptic terminal^9–169–12, 14–16^. For the presynaptic protein aSYN the concept of biomolecular condensates opens new avenues for understanding its role at the synapse, since, besides its clear link to disease ^17, 78, 89^, its normal physiological function remains elusive. To date, several reports show that aSYN can undergo droplet formation ^19, 23– 25, 90^ and that these condensates show hardening, which is potentially relevant for the transition to pathological states ^21, 22^. Here, we demonstrate that the phase separation of aSYN can be regulated via C-terminal interaction. We find that the presence of spermine and also Ca^2+^ promotes fast droplet formation of aSYN under otherwise same conditions. This aligns with recent studies demonstrating a facilitatory effect of Ca^2+^ and Mn^2+^ on aSYN phase separation ^24, 25^. However, we do not observe facilitation of aSYN droplet formation at physiologically relevant Ca^2+^ concentrations estimated to be 200-300 µM Ca^2+^ or even below during synaptic stimulation ^62, 63^. Therefore we speculate that this reflects an electrostatic modulation, rather than a direct regulatory role of Ca^2+^ itself. We demonstrate, that in cells, co-expression of aSYN with an interaction partner, namely VAMP2 leads to the formation of aSYN condensates. aSYN has been shown before to interact with VAMP2, in particular via its C-terminal region ^49–51^, however, a role for the induction of aSYN phase separation is new. Our findings on the role of VAMP2 on aSYN phase separation might reflect earlier reports which indicate that aSYN dispersion upon synaptic activity is inhibited when SNARE-mediated SV exocytosis is blocked ^83, 91^. These experiments were performed under normal Ca^2+^ conditions and therefore emphasize that not Ca^2+^ entry but rather vesicle exocytosis and the respective conformational state of the individual SNARE proteins can influence the spatiotemporal organization of aSYN, congruent with our findings here.

In addition to the role of VAMP2 on aSYN phase separation, our results demonstrate that the formation of aSYN condensates in cells is dependent on aSYN’s binding to lipid membranes. Here, the aSYN A30P variant with decreased lipid binding completely abolished condensate formation. This indicates that aSYN forms membrane-associated condensates rather than undergoing droplet formation in solution, which is in accordance with its well-described property of vesicle binding ^29–37^. Interestingly, the A30P variant abolishes aSYN condensate formation completely, while in vitro aSYN binding is only reduced by about half ^80, 92^. Again, this reflects previous findings, where aSYN A30P completely abolishes the membrane accumulation of aSYN in yeast and at the synapse ^82, 83^. Further to that, we show that aSYN condensates in cells have a regulatory role on clustering vesicles. In contrast to the vesicle clusters which have been found upon synaptophysin/synapsin co-expression ^84^, we find that vesicles are less regular in size, which might reflect a differential function of aSYN in SV clustering. In this context, different vesicle cluster entities formed by distinct condensates have been demonstrated lately ^85^. Recent findings in the lamprey giant reticulospinal synapse verify a role of aSYN in SV clustering directly at the synapse ^93^, supporting our findings and previous in vitro experiments on aSYN vesicle clustering ^53–55^. In the lamprey synapse, aSYN depletion not only effected the distal pool of vesicles ^93^, as has been found upon synapsin depletion or interference with synapsin phase separation ^10, 94^ but also the proximal vesicle pool located adjacent to the active zone, emphasizing different and/or supplementary roles for aSYN and synapsin ^26, 93^.

The interaction of VAMP2 and aSYN involved in aSYN phase separation is mediated by the JM-domain of VAMP2. This is interesting since immunoprecipitation experiments show that the N-terminal proline-rich domain of VAMP2 mediates VAMP2/aSYN interaction ^49^. Our experiments do not exclude an interaction between the N-terminal domain of VAMP2 and aSYN’s C-terminal region but rather indicate that another region is responsible for the induction of aSYN phase separation. The role of the JM-domain was surprising, first because proline-rich domains are known to mediate protein interactions and phase separation and second because other proteins, i.e. syntaxin-1A, also exhibit similar positively charged motifs within their stop transfer signal. The role for VAMP2 was specific, but we do not exclude the possibility that other VAMP family members might have similar effects on aSYN, which is indeed likely since aSYN overexpression has been shown to interfere with other vesicle transport mechanisms, i.e. ER-Golgi transport ^95, 96^. Sterically the JM-domain of VAMP2 is in close proximity to the lipid membrane as well as the aSYN residues, which have been demonstrated to interact most strongly with VAMP2 ^50^. An additional regulatory role for the proline-rich N-terminal region of VAMP2 and more C-terminal residues of aSYN, i.e. in attracting further protein partners or via S129 phosphorylation is to be explored.

In summary, our results demonstrate that aSYN phase separation is regulated via its C-terminal domain. This can be achieved via electrostatic interactions between the negatively charged C-terminus and positively charged molecules, such as spermine or Ca^2+^, or via protein-protein interaction with VAMP2, in particular its JM-domain. While Ca^2+^ seems to modulate aSYN phase separation only at high concentrations, the JMD peptide allowed induction of aSYN phase separation at a concentration that aligns with the estimated VAMP2 concentration at the synapse, which is reported to be 170 μM ^97^. While we demonstrate that aSYN phase separation is linked to VAMP2 and therefore could be influenced during SNARE complex formation and exocytosis, this does not preclude a potential function for aSYN during SV endocytosis as suggested by previous studies ^98–101^. Our findings delineate a molecular mechanism for the regulation of aSYN phase separation which will allow us to further explore its role during vesicle cycling.

## Acknowledgment

J.L. acknowledges funding from the Royal Society (Royal Society Dorothy Hodgkin Research Fellowship, DHF/R1/201228), from the Addenbrooke’s Charitable Trust (Grant Award 900325), the Leverhulme Trust (Research Project Grant, RPG-2022-257), as well as a Career Support Fund from the University of Cambridge. We thank Maria Guerra Martin and Jaime Llodra Gonzalez for EM supports. AJW acknowledges funding from Blood Cancer UK (21002) and the UK Medical Research Council (MR/T012412/1). Furthermore, we thank Dr David Gershlick for discussion on the manuscript draft.

## Author Contributions

Conceptualization, J.L.; Methodology, A.A., C.H., N.M., and J.L.; Investigation, A.A., F.R., C.H., K.S., N.M., and J.L; Writing – Original Draft, J.L. and A.A.; Writing – Review & Editing, J.L., A.A, C.H., and A.J.W.; Funding Acquisition, J.L.; Supervision, A.J.W., and J.L.

## STAR METHODS

### RESOURCE AVAILABILITY

#### Lead contact

Further information and requests for resources and reagents should be directed to and will be fulfilled by the lead contact, Janin Lautenschlaeger.

#### Materials availability

Plasmids generated in this study are available from the lead contact with a completed material transfer agreement.

#### Data and code availability

Data reported in this paper will be shared by the lead contact upon request. Any additional information required to reanalyse the data reported in this paper is available from the lead contact upon request.

### EXPERIMENTAL MODEL AND STUDY PARTICIPANT DETAILS

#### Cell culture and transfection

HeLa cells were obtained from the European Collection of Cell Cultures (ECACC 93021013) and grown in Dulbecco’s modified Eagle’s Medium (DMEM) high glucose (31966-021, Gibco) supplemented with 10% fetal bovine serum (FBS, F7524, Sigma) and 1% Penicillin/Streptomycin (P0781, Sigma). Cells were grown at 37 °C in a humidified incubator with 5% CO_2_. Cells were tested for mycoplasma contamination using MycoStrip^TM^ (IvivoGen, Toulouse, France). Cells were plated at 20 000 cells/well in 48-well plates (Cellstar, 677 180, Greiner bio-one) for incuCyte experiments or in 8-well ibidi dishes (80807, ibidi, Gräfelfing, Germany) for live cell confocal imaging. Cells were transfected using Fugene HD Transfection reagent according to the manufacturer’s protocol (E2311, Promega). Briefly, per reaction 12.5 μL OptiMEM (31985-062, Gibco) were set up in 1.5 mL sterile Eppendorf tubes. A total of 250 ng of DNA and 0.75 μL of Fugene reagent were added and incubated for 15 min at room temperature. The transfection mix was added onto the cells for 1 min and then topped up with 300 μL complete media.

### METHOD DETAILS

#### Protein expression and purification of aSYN

Recombinant wild-type human full-length aSYN cloned in vector pET28a (Addgene #178032) was transformed into BL21(DE3) competent Escherichia coli (C2527, NEB, Ipswich, US). Bacteria were cultured in LB media supplemented with 50 μg/mL kanamycin (37 °C, constant shaking at 250 rpm). Expression was induced at an OD600 of 0.8 using 1mM isopropyl β-D-1-thiogalactopyranoside (IPTG) and cultured overnight at 25 °C. Cell pellets were harvested by centrifugation at 4000 g for 30 minutes (AVANTI J-26, Beckman Coulter, USA). aSYN was purified using a protocol previously described ^102^. Briefly, the cell pellet was resuspended in lysis buffer (10 mM Tris, 1 mM EDTA, Roche cOmplete EDTA free protease inhibitor cocktail, pH 8). The cells were disrupted using a cell disruptor (Constant Systems, Daventry, UK) and were ultracentrifuged at 4 °C, 40 000 rpm for 20 minutes (Ti-45 rotor, Optima XPN 90, Beckman Coulter, USA). The supernatant was collected and heated for 20 minutes at 70 °C to precipitate heat-sensitive proteins, followed by ultracentrifugation as above. Streptomycin sulfate (5711, EMD Millipore, Darmstadt, Germany) was added at a final concentration of 10 mg/mL to the supernatant and continuously stirred at 4 °C for 15 minutes to precipitate DNA, followed by ultracentrifugation as above. Ammonium sulfate (434380010, Thermo Scientific) was added at a final concentration of 360 mg/mL to the supernatant and continuously stirred at 4 °C for 30 minutes to precipitate the protein. The precipitated protein was then centrifuged at 500 g for 15 min, dissolved in 25 mM Tris, pH 7.7, and dialyzed overnight against the same buffer to remove salts. The protein was purified using ion exchange on a HiTrap^TM^Q HP 5mL anion exchange column (17115401, Cytiva, Sweden) using gradient elution with 0-1M NaCl in 25 mM Tris, pH 7.7. The collected protein fractions were run on SDS-PAGE and pooled fractions were further purified using size-exclusion chromatography on a HiLoadTM 16/600 SuperdexTM 75 pg column (28989333, Cytiva, Sweden). The fractions were collected, and their purity was confirmed using SDS-PAGE analysis. Protein concentrations were determined by measuring absorbance at 280 nm using an extinction coefficient of 5,600 M^−1^cm^−1^. The monomeric protein was frozen in liquid nitrogen and stored in 25 mM HEPES buffer pH 7.4 at -70 °C. pET28a Cdk2ap1CAN was a gift from Lin He (Addgene plasmid # 178032; http://n2t.net/addgene:178032; RRID:Addgene_178032) ^103^.

#### Protein expression and purification of VAMP1-96 and syntaxin1-265

Recombinant wild-type human VAMP1-96 and syntaxin1-265 cloned in vector pOPINS and pET28a (Addgene #66711, #178032) respectively were transformed into BL21(DE3) competent Escherichia coli (C2527, NEB, Ipswich, US). Bacteria were cultured in LB media supplemented with 50 μg/ml kanamycin (37 °C, constant shaking at 250rpm). Expression was induced at an OD600 of 0.8 using 1mM isopropyl β-D-1-thiogalactopyranoside (IPTG) and cultured overnight at 25 °C. Cell pellets were harvested by centrifugation at 4000 g for 30 minutes (AVANTI J-26, Beckman Coulter, USA). HisSUMO-tagged VAMP1-96 was purified using Ni-NTA chromatography using the following protocol. Briefly, the cell pellet was resuspended in lysis buffer (25 mM HEPES, 300 mM NaCl, Roche cOmplete EDTA free protease inhibitor cocktail, pH 7). The cells were disrupted using a cell disruptor (Constant Systems, Daventry, UK) and were ultracentrifuged at 4 °C, 40 000 rpm for 20 minutes (Ti-45 rotor, Optima XPN 90, Beckman Coulter, USA). The supernatant was incubated with Ni-NTA resin overnight at 4 °C which was then loaded on the column. The column was washed with 25 mM imidazole and the His-tagged protein was eluted with 250 mM imidazole, pH 7. The eluted protein was incubated with SUMO protease (10 unit/mg of protein, SAE0067, Sigma) and was set for overnight dialysis (25 mM HEPES, 300 mM NaCl, pH 7) at 4 °C. SUMO protease and uncleaved protein were removed incubating the dialyzed protein solution with Ni-NTA resin for 2 hours at 4 °C. The flow-through with His-cleaved VAMP1-96 protein was collected and its purity was confirmed using SDS-PAGE analysis. Protein concentrations were determined by measuring absorbance at 280 nm using an extinction coefficient of 13,980 M^−1^cm^−1^. The monomeric protein was frozen in liquid nitrogen and stored at -70 °C. His-tagged Syntaxin1-265 was purified using Ni-NTA chromatography as above. Briefly, the cell pellet was resuspended in lysis buffer (25 mM HEPES, 300 mM NaCl, 1 mM DTT, Roche cOmplete EDTA free protease inhibitor cocktail, pH 7.4). The supernatant obtained after cell disruption and centrifugation was directly loaded on to the Ni-NTA column. The column was washed with 25 mM imidazole and the His-tagged protein was eluted with 250 mM imidazole and was dialyzed against buffer (25 mM HEPES, 300 mM NaCl, 1 mM DTT, pH 7.4) overnight. The protein was further purified using size-exclusion chromatography on a HiLoadTM 16/600 SuperdexTM 200 pg column (28989335, Cytiva, Sweden). The fractions were collected, and their purity was confirmed using SDS-PAGE analysis. Protein concentrations were determined by measuring absorbance at 280 nm using an extinction coefficient of 7450 M^−1^cm^−1^. The monomeric protein was frozen in liquid nitrogen and stored at -70 °C. pOPINS-UBE3C was a gift from David Komander (Addgene plasmid # 66711; http://n2t.net/addgene:66711; RRID:Addgene_66711) ^104^. pET28a Cdk2ap1CAN was a gift from Lin He (Addgene plasmid # 178032; http://n2t.net/addgene:178032; RRID:Addgene_178032) ^103^.

#### Labeling of proteins

Labeling of proteins was performed in bicarbonate buffer (C3041, Sigma) at pH 8 using NHS-ester active fluorescent dyes. VAMP1-96 and syntaxin1-256 were labeled using Janelia Fluor 549 SE (6147, Tocris), aSYN was labeled using AlexaFluor 488 5-SDP ester (A30052, Invitrogen Thermo Fisher). Excess-free dye was removed by buffer exchange using PD10 desalting columns (IP-0107-Z050.0-001, emp BIOTECH, Generon). Labeled protein concentrations were estimated using molar extinction coefficients of the dyes, ε_555 nm_ = 101, 000 M^−1^cm^−1^ for Janelia Fluor 549 SE; ε_494 nm_ = 72,000 M^−1^cm^−1^ for Alexa-488 5-SDP ester.

#### Phase separation assays including turbidity measurements, confocal imaging, and sedimentation-based assays

Phase separation assays were performed in 25 mM HEPES, pH 7.4 unless mentioned otherwise. Phase separation was induced by mixing aSYN and PEG 8000 (BP223, Fisher Bioreagent) in the presence or absence of calcium (21108, Sigma), spermine (S2876, Sigma), or VAMP2 peptides (Custom synthesis with Proteogenix, Schiltigheim, France) as indicated respectively. **Turbidity measurements** – Phase-separated samples were set up as described above. The turbidity of the samples was measured at 350 nm, 25 °C using 96-well Greiner optical bottom plates on a CLARIOstar plate reader (BMG LABTECH, Ortenberg, Germany) under quiescent conditions. A sample volume of 100 μL was used, and readings were taken within 5 minutes of sample preparation. For phase diagrams, the raw turbidity data are plotted with background subtraction. Data were obtained from at least three independent sets of biological samples and were plotted using OriginPro 2018. **Confocal Microscopy**-images for phase-separated samples were acquired on an LSM780 confocal microscope (Zeiss, Oberkochen, Germany) using a 63x oil immersion objective. Images were taken in brightfield mode and/or using aSYN supplemented with 1% Alexa 488 labeled aSYN. For colocalization experiments, aSYN phase separation was induced in the presence of 2 mM Ca^2+^ and 15% PEG 8000. aSYN was supplemented with 1% Alexa 488 labeled aSYN. VAMP1-96 and syntaxin1-265, supplemented with 1% Alexa 594 labeled protein, were added after droplet formation and imaged after 30 minutes of incubation. Data were obtained using the same imaging conditions for VAMP1-96 and syntaxin, respectively. Images analysis was performed in FIJI ^105^. Briefly, a mask for aSYN droplets was generated in the 488 channel which was used to measure the intensity of aSYN in the 488 channel and the intensity of VAMP1-96 and syntaxin1-265 in the 594 channel. The ratio of VAMP1-96 and syntaxin1-265 to aSYN intensity was plotted. Data from at least three independent sets of biological samples were obtained. **Sedimentation-based assays** – aSYN phase separation was induced in the presence of 2 mM Ca^2+^ and 15% PEG 8000, followed by the addition of VAMP1-96 protein. 50 μL reactions were set up and incubated for 20 minutes. Samples were centrifuged at 25,000 × g for 20 minutes at 25 °C to separate the dense phase (pellet fraction) and the light phase (supernatant). The supernatant was carefully removed, and the pellet was resuspended in 8 M urea. The samples were run on a 15% SDS-PAGE gel and were visualized using Coomassie blue staining (Quick Coomassie stain, Protein Ark). The gels were scanned on a ChemiDoc^TM^MP Imaging System (BioRAD, UK) and the saturation concentration (C_sat_) was calculated from the band intensity referenced to a known aSYN standard using FIJI software^105, 106^.

#### Constructs / Plasmids and transfection

Wild-type human full-length aSYN and VAMP2 were cloned from cDNA obtained from human neuroblastoma cells (SH-SY5Y) and cloned into the pEYFP-N1 and pMD2.G vector (Addgene #96808, #12259) respectively. Synaptojanin 145 and 170, endophilin-A1, and Hsc70 constructs were purchased from Addgene (#22291, #22292, #47403, #86031), pcDNA3-FLAG-Synaptojanin 1-145 and Synaptojanin 1-170 were a gift from Pietro De Camilli (Addgene plasmid # 22291; http://n2t.net/addgene:22291; RRID: Addgene_22291; Addgene plasmid # 22292; http://n2t.net/addgene:22292; RRID: Addgene_22292) ^107^, Endophilin Full Length was a gift from Peter McPherson (Addgene plasmid # 47403; http://n2t.net/addgene:47403; RRID: Addgene_47403) ^108^, pCDNAZeo(-)HSC73AS was a gift from Janice Blum (Addgene plasmid # 86031; http://n2t.net/addgene:86031; RRID: Addgene_86031) ^109^. All other synaptic proteins, synphilin (Sph1), Rab-3A, rabphilin-3A, VPS35, auxilin, syntaxin-1A, and SNAP25 were cloned from SH-SY5Y cDNA and were then inserted into the pMD2.G vector. Synaptotagmin-1 was cloned from SH-SY5Y cDNA into the pCDNA3.1 vector with an N-terminal mScarlet tag (Addgene #16015, Addgene #85045). Gibson assembly was performed upon PCR amplification (Q5 Hot start HiFi 2xMM, M0494S, 2xHiFi DNA Assembly MM, E2621S, NEB, Ipswich, US). aSYN 96AAA and A30P were generated using KLD substitution (M0554S, NEB, Ipswich, US). All sequences were verified by sequencing. 5HT6-YFP-Inpp5e was a gift from Takanari Inoue (Addgene plasmid # 96808; http://n2t.net/addgene:96808; RRID:Addgene_96808) ^111^. pMD2.G was a gift from Didier Trono (Addgene plasmid # 12259; http://n2t.net/addgene:12259; RRID:Addgene_12259). pcDNA3.1/V5-His Snk/Plk2 was a gift from Wafik El-Deiry (Addgene plasmid # 16015 ; http://n2t.net/addgene:16015 ; RRID:Addgene_16015) ^113^. pmScarlet_Giantin_C1 was a gift from Dorus Gadella (Addgene plasmid # 85048 ; http://n2t.net/addgene:85048 ; RRID:Addgene_85048) ^115^.

#### Confocal microscopy, IncuCyte, and image analysis

Cells in 48-well plates were imaged with the IncuCyte S3 (Essen BioScience, Newark, UK). Phase brightfield and green fluorescence images were taken using a 20x objective at a 4-hour interval at 200 ms exposure, condensate formation (% of cells showing condensate formation) was evaluated 16 hours after transfection. At least three biological repeats with three technical repeats each were analyzed blinded to the investigator. Live cell confocal imaging was performed on an LSM780 microscope (Zeiss, Oberkochen, Germany) using a 63x oil immersion objective. YFP fluorescence was excited with the 514 laser at 2% laser power, mScarlet was excited using the 561 laser at 2% laser intensity. For fluorescence recovery after photobleaching (FRAP) experiments images were taken with the 63x oil immersion objective, 20x zoom, 128x128 pixel resolution, at an imaging speed of 60 ms/image. Three pre-bleach images were acquired before the ROI was bleached with 100 iterations at 100% laser power using the 514 laser. Fluorescence recovery was recorded for 100 cycles. FRAP analysis was performed in FIJI using the FRAP profiler v2 plugin (Hardin lab, https://worms.zoology.wisc.edu/research/4d/4d.html). Small aliphatic alcohols were used to evaluate the role of hydrophobic interaction for aSYN condensate formation. 1,6-hexanediol (240117, Sigma, USA) and 1,3-propanediol (P50404, Sigma, China) were prepared as 6% stock solutions in complete DMEM media and were added to the cells in a 1:1 ratio after the first image was taken. Condensate size, number of condensates per cell, and the ratio of condensate intensity and aSYN cytosolic intensity for aSYN WT and aSYN 96AAA were analyzed using FIJI ^105^. Analysis was performed blinded to the investigator. Pearson Correlation Coefficients were calculated using the ColocFinder Plugin (https://imagej.nih.gov/ij/plugins/colocalization-finder.html). Profile plots were generated in FIJI ^105^ and plotted in Prism.

#### NMR spectroscopy

Recombinant [^15^N]-labeled wild-type human full-length aSYN was grown in M9 minimal medium supplemented with a vitamin mix, trace elements and the appropriate nitrogen (^15^NH_4_) and carbon (^12^C-glucose) sources and purified as described above. [^15^N]-labelled aSYN was dialysed into 20 mM phosphate buffer, pH 7.0. 200 μL samples were prepared for NMR spectroscopy containing 40 μM aSYN WT, the respective concentration of peptide (Custom synthesis by Proteogenix, Schiltigheim, France) and 10% ^2^H_2_O (Thermo Scientific Chemicals). To probe the interaction of peptide with aSYN at a residue specific level, we used multi-dimensional NMR spectroscopy and recorded ^1^H-^15^N-BEST-TROSY spectra atitrating the concentration of the JMD long peptide (50, 100, 150 μM peptide) at a fixed concentration of aSYN of 40 μM. The inter-scan delay was set to 0.3 s. NMR experiments were carried out at 800MHz and 600MHz using Bruker Avance spectrometers equipped with 5 mm triple resonance inverse cryoprobes, at 298 K. Assignments of the resonances in the ^1^H-^15^N-BEST-TROSY spectra of aSYN were derived from previous studies ^80^. Spectra were internally referenced to the ^1^H_2_O signal at 4.70 ppm, processed with TopSpin version 2.1 (Bruker), and analysed with SPARKY (Goddard TD & Kneller DG, SPARKY 3, University of California, San Francisco).

#### Correlated Light and Electron Microscopy

HeLa cells were cultured on gridded sapphire discs with carbon coating (K950X Turbo Evaporator, Quorum Tecknologies, East Sussex UK; Leica EM-ACE200 Vacuum Coater, Vienna, Austria), transfected as above using co-expression of aSYN YFP and VAMP2, and imaged the next day on an LSM780 confocal microscope (Zeiss, Oberkochen, Germany) using a 40x oil immersion objective. The cells were then proceeded to high-pressure freezing and freeze substitution ^110, 112, 114^ using a Leica EM-HPM100 and EM-AFS2/FSP (Leica Microsystems, Vienna Austria). Briefly, the specimens were dealt in 0.1% tannic acid in anhydrous acetone at -90 °C for at least 24 hrs, washed with pure acetone at -90 °C, and substituted in 1% osmium tetroxide, 0.1% uranyl acetate, and 1% pure water in acetone, with temperature control at -90 °C for 72 hrs, -60 °C for 8 hrs, and -30 °C for 8 hrs and interval of 30 °C/hr. After washing with pure acetone at 4 °C, the cells were embedded in Epon and Araldite mixture (TAAB Laboratories Equipment Ltd., Reading UK). After polymerization at 65 °C for a few days, ultrathin-sections (∼60 nm) obtained by Ultramicrotome (Leica EM-UC7/Artos-3D) were mounted on formvar-carbon films/EM grids, stained with lead citrate and uranyl acetate, and then observed by 200kV transmission electron microscope (FEI Talos, Oregon USA) with Ceta-16M CMOS-based camera (4kx4k pixels under 16-bit dynamic range). Imaging was carried out within the correlated region of preceding confocal fluorescence imaging.

### QUANTIFICATION AND STATISTICAL ANALYSIS

All data are represented as mean ± standard error (SD). Statistical parameters are reported in the Figures and the corresponding Figure Legends. Statistical analysis was performed in GraphPad Prism 9.3.1 using unpaired two-tailed t-test or one-way ANOVA with Dunnett’s multiple comparison test. For 1,6-hexanediol and 1,3-propanediol experiments paired two-tailed t-test and RM one-way ANOVA with Geisser-Greenhouse correction and Dunnett’s multiple comparisons test were performed, respectively. P-values of less than 0.05 were considered significant with * p < 0.05; ** p < 0.01; *** p < 0.001; and **** p < 0.0001.

---

**Figure S1.**
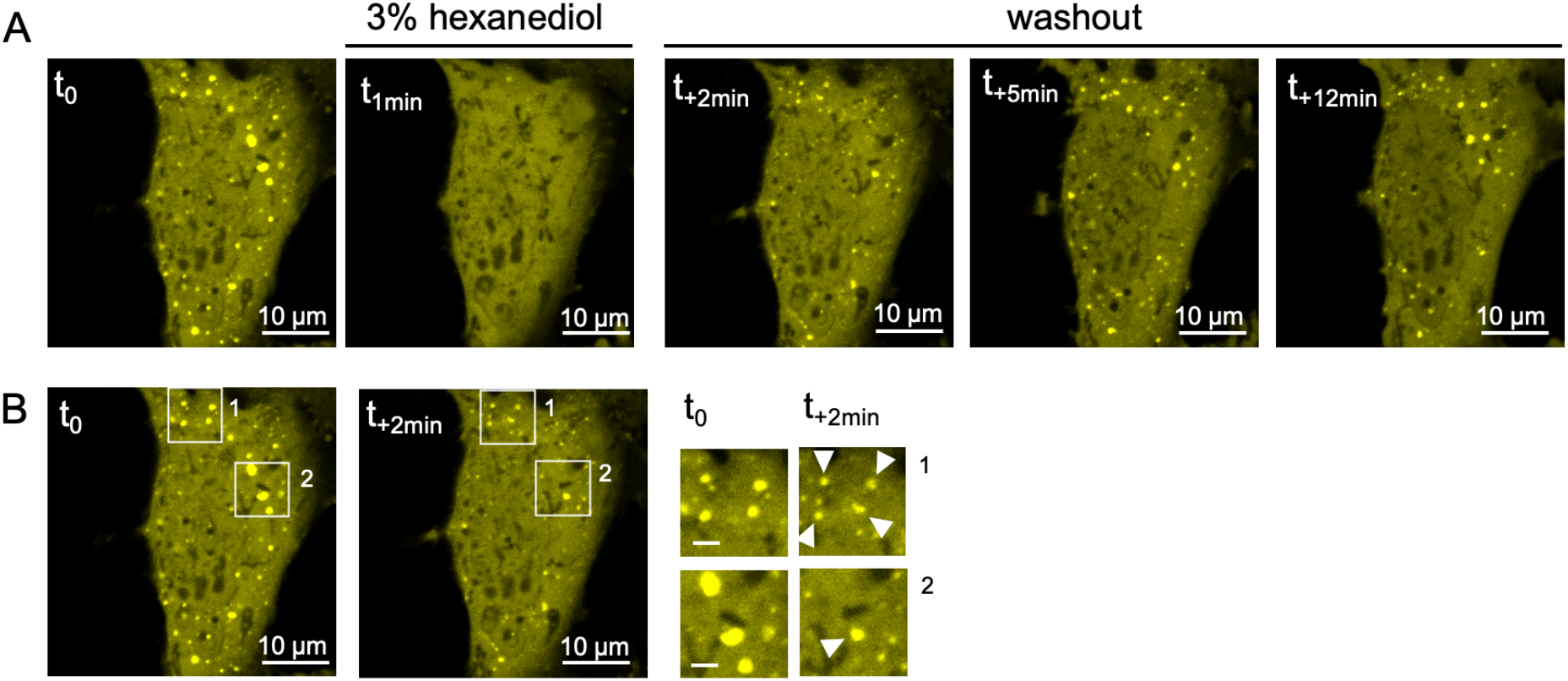
Sensitivity of aSYN condensates to 1,6 hexanediol. (A) aSYN YFP condensates show dispersal upon incubation with 3% 1,6 hexanediol with fast recovery after 1,6 hexanediol washout. (B) Reassembly of aSYN condensates at the same regions (arrowheads).

**Figure S2.**
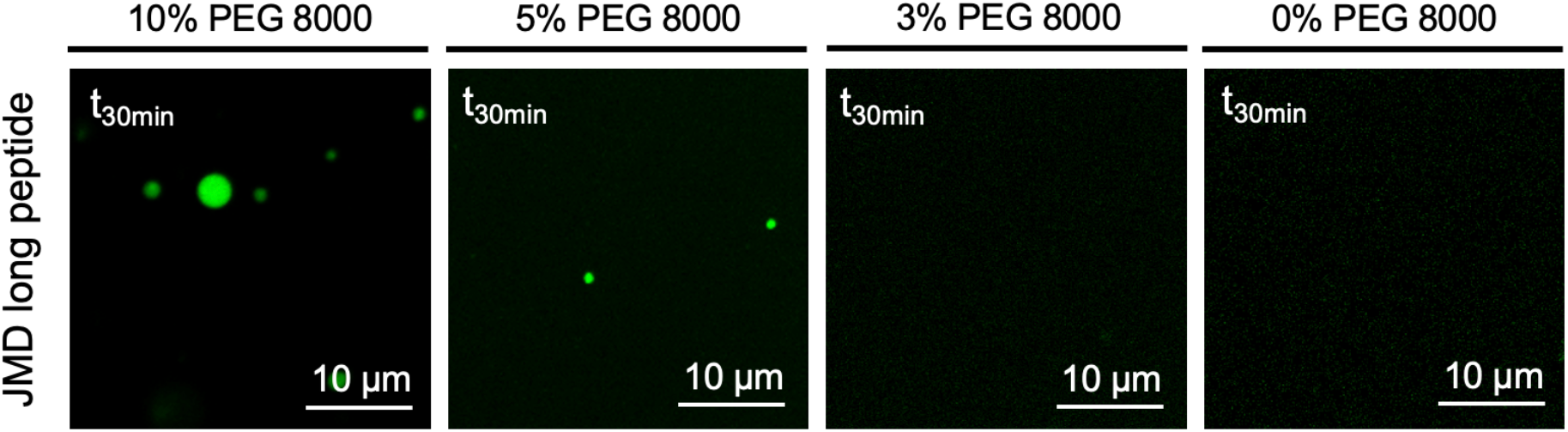
aSYN droplet formation at low PEG concentrations. (A) aSYN droplet formation in the presence of 150 µM JMD long peptide, PEG concentrations as indicated, aSYN concentration used: 40 µM

**Figure S3.**
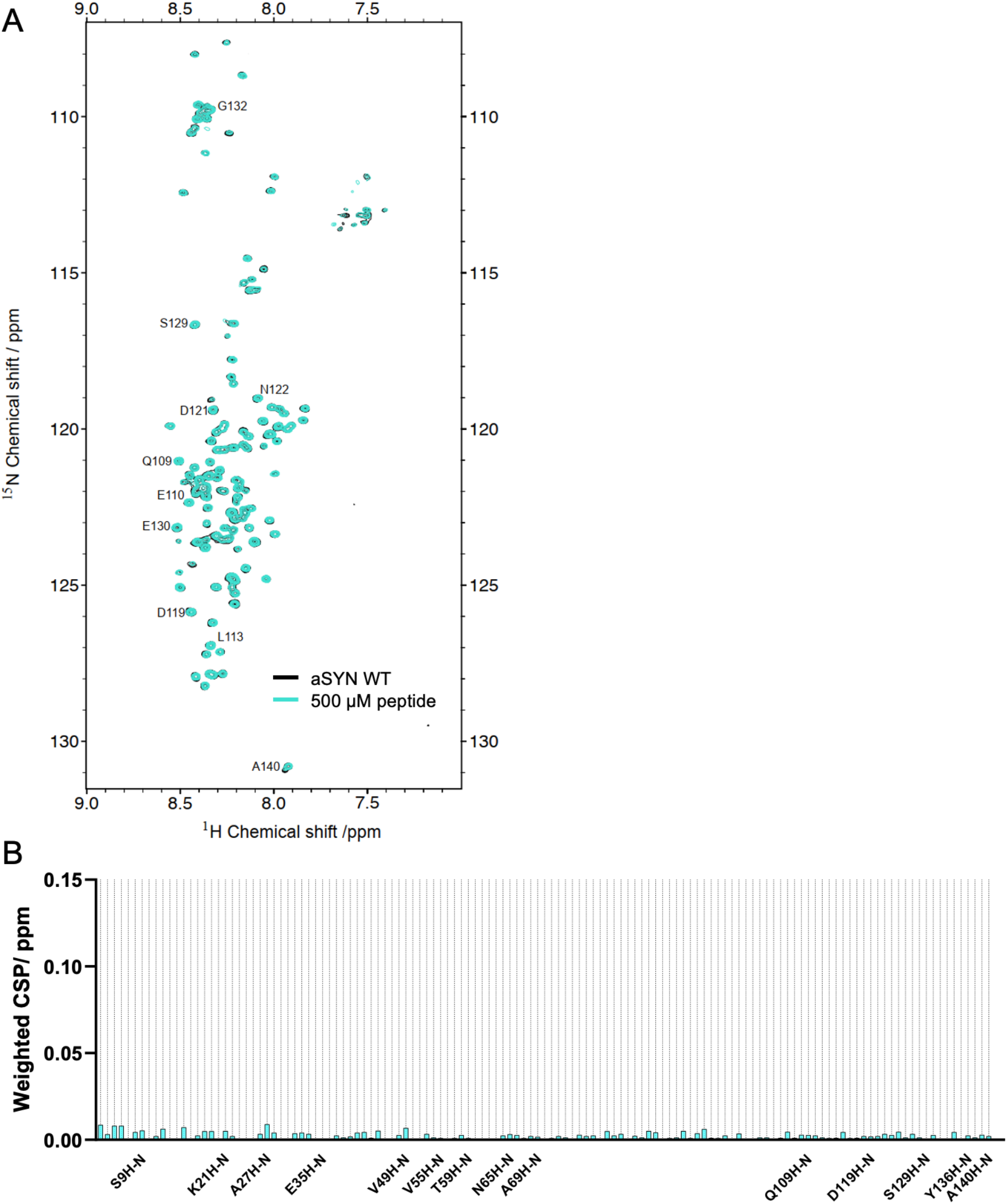
^1^H-^15^N-BEST-TROSY spectra for NT long peptide 1. (A) Overlappped ^1^H-^15^N-BEST-TROSY spectra of aSYN without (black) and in the presence of NT long peptide 1 (turquoise). aSYN concentration used: 40 µM, peptide concentration: 500 µM. (B) Weighted average (of ^15^N and ^1^H) chemical shift perturbation 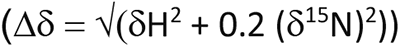 of residues in aSYN in the presence of 500 μM NT long peptide 1.

